# Life-stage-specific specialities in the cell atlases of the *Clytia hemisphaerica* planula and medusa

**DOI:** 10.64898/2026.02.16.705741

**Authors:** Anna Ferraioli, Julia Ramon Mateu, Marc Meynadier, Thomas Lamonerie, Sophie Pagnotta, Sandra Chevalier, Marta Iglesias, Sebastian R Najle, Arnau Sebé-Pedrós, Marie-Jeanne Arguel, Julie Cazareth, Virginie Magnone, Evelyn Houliston, Richard R. Copley

## Abstract

Jellyfish have complex life-cycles, but there has been limited exploration of how this is achieved at the cellular level. We used single-cell transcriptomics to assemble a cell atlas for the planula larva of *Clytia hemisphaerica*, and compared it to an updated cell atlas for the medusa (jellyfish) stage. The cells of the planula fell into the same broad categories as for the medusa: ectoderm, gastroderm, interstitial cells (i-cells), nematocytes (stinging cells), neurons and secretory cells. Although the planula cells generally showed less diversity than medusae within each category, cells with specialized features unique to their stage could be distinguished by their transcriptional profiles as well as by ultrastructure. Some planula-specific types were identified: aboral secretory cells involved in settlement, and a cell type attributed a role in immunity or post-metamorphic theca production. Distinct transcriptome profiles within different regions of the ciliated planula ectoderm reflected different post-metamorphosis fates of domains along the oral-aboral axis. Inspection of the cell clusters showing significant similarity of marker genes between planula and medusa, and inference of similarity using a statistical model of marker gene presence/absence, revealed correspondences between families of cells from planula and medusa rather than precise cell identities.

## Introduction

In animals with complex life-cycles the same genome directs cells to develop into distinct autonomous body plans, but the degree to which the cells themselves may be the same or different between stages is a question that has not been widely addressed. The constraints of metazoan ancestry suggest that different life-cycle stages will use similar cell types to make nervous systems, muscles and so on, but these are likely to require some degree of specialisation dependent on the niche occupied. Equally, stages may require cells that are not required in other stages, for example, digestive cells in feeding adults, absent in non-feeding larvae.

Among marine species where these questions have been examined, larval cell types of the anthozoan cnidarian *Nematostella* have been compared to the adult polyps, revealing larval-specific cell types associated with the apical organ, but ‘remarkably related’ transcriptional programs for larval and adult differentiated cell types, indicating limited rearrangement during metamorphosis [1]. The *Nematostella* ‘planula’ larva, however, transitions fairly smoothly into an adult polyp [2], and does not undergo the abrupt settlement characteristic of cnidarians from other anthozoan and medusozoan clades. In contrast to *Nematostella*, the sponge larval cell types of *Amphimedon queenslandica*, which does settle, were described as showing ‘remarkable differences compared with adult cell types’, and only one cell type, archaeocytes, equivalent between the two stages [3]. Given that archaeocytes are pluripotent [4], it is difficult to regard them as a shared differentiated cell type. The enteropneust hemichordate *Schizocardium californicum*, which metamorphoses abruptly, shows a striking discordance between larval and adult cell types [5]. Arnone and co-workers have investigated similarities between the larva and adult stages of the sea urchin *Paracentrotus lividus*, identifying developmentally conserved and diverse units in the post-metamorphic animal, including muscle, but no direct similarity between larval and juvenile neurons [6].

We have previously produced a cell atlas for the medusa stage of the cnidarian *Clytia hemisphaerica* [7]. *Clytia* has a canonical hydrozoan life cycle with three stages: planula larvae, benthic colonies of connected polyps, and swimming medusae (jellyfish). Cnidarian planula larvae are ciliated, with a similarity to acoelomorph flatworms that has led to the proposal that the cnidarian-bilaterian ancestor may have shared this form [8], but previous work has shown that in *Clytia* the medusa stage is more bilaterian-like in its gene deployment [9]. To enquire farther into the cellular complexity of the planula and medusa, we generate a cell atlas for the planula larva and compare it to an updated medusa cell-type atlas. We identify broad conserved cell-type categories that are shared by larvae and medusa, as well as cell types that are specific to one or the other stage.

## Results

### An updated medusa cell atlas

We generated two new scRNA-seq datasets for the *Clytia hemisphaerica* medusa, mapping these, and data from [7] to a chromosome-level genome assembly (see methods). Analysis using a standard scanpy pipeline [10], with Harmony integration between datasets [11] produced 22 cell clusters, with a total of 30,544 cells. Our analysis showed prominent batch effects between datasets. To counter this, we created a pseudobulk dataset with each batch as a replicate, and used DESeq2 to calculate statistically significant marker genes for each post-Harmony cluster, allowing for the batch in the experimental design (see methods). We calculated the statistical significance of cell type marker gene list overlap between all pairs of medusa cell types (see methods). Clustering based on the significance of the similarity of overlap revealed subtypes within broad categories of gastroderm, ectoderm, nematocytes, neurons, gland cells and stem cells/germ cells, consistent with our previous analyses (**Fig. 1a and 1b**) [7]. Some classes, including neurons, can be further split into distinct subtypes at higher clustering resolutions. We have previously considered medusa neurons sub-types in more detail [7]; others distinctions e.g. velum and subumbrella striated muscle, although giving separable molecular signatures at high resolution, are not pursued here.

**Figure 1:**
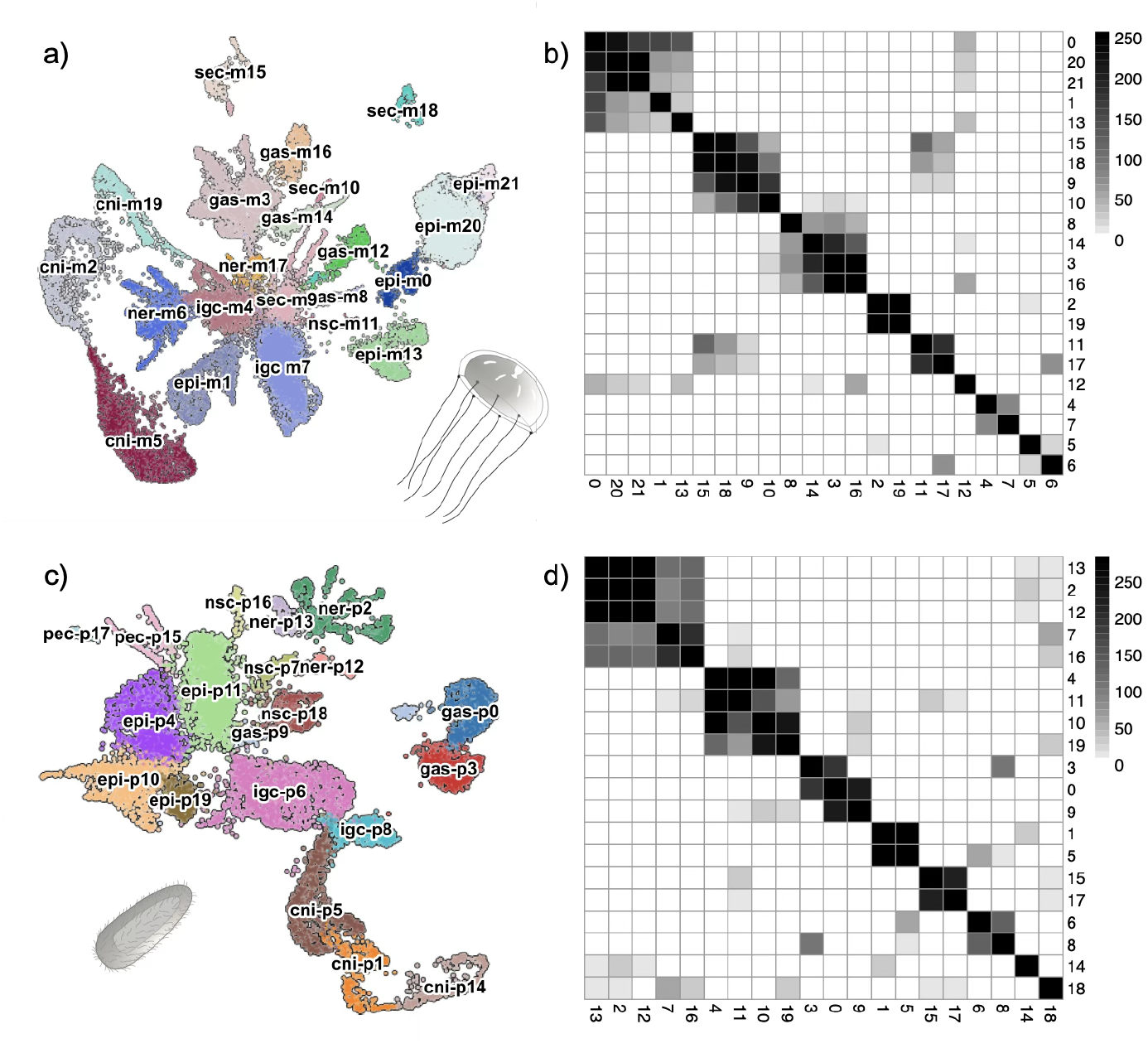
*Clytia* single-cell dataset representations. **a**,**c)** UMAPs colored by Leiden cell cluster, with schematic diagrams of *Clytia* medusa and planula (aboral end down) larva stage; **b)** self-similiarity of inferred Leiden cluster marker gene sets for medusa **d)** and planula single cell data sets (see methods). In the UMAPs Leiden cluster numbers are prefixed with ‘m’ for medusa’ and ‘p’ for planula and these are then prefixed with cell category abbreviations: cni, nematocytes; epi, epidermis; gas, gastrodermis; gla, gland; igc i-cells/germ cells; ner, neuron; nsc, neurosecretory; sec, secretory.

### Planula cells: annotation and atlas

We generated four experimentally distinct scRNA-seq datasets for planula larvae, processing them in the same way as the medusa data. Three datasets were from planulae collected on the second day after fertilisation, and one from the third day. For planula cells, we found that fixation immediately after dissociation was essential to preserve mRNA quality (see methods). The combined dataset yielded 13,655 cells in 20 clusters **(Fig. 1c & 1d)**. As with the medusa, to see higher-level similarities, we calculated the significance of marker gene overlap between planula clusters. To facilitate the transfer of annotation from medusa cell clusters to planula, we calculated the statistical significance of cell type marker gene list overlap between all pairs of cell types (**Fig. 2** - see methods). Using previously assigned medusa maker genes, we were able to assign the clusters to the same broad medusa cell categories, namely gastroderm, ectoderm, i-cells, nematocytes, neural and secretory. The statistical analysis led us to classify the mucous cells and ‘granular’ cells of [12] within broader classes (see below). *In situ* hybridization performed on staged planulae with selected markers allowed us to consolidate these assignments and to propose individual cluster identities (**Fig. 3 & S2**). Transmission Electron Microscopy (TEM) of planulae fixed 2 days after fertilisation (‘2-day planulae’) showed cellular features reflecting the transcriptional features of the clusters **(Fig. 4)**.

**Figure 2:**
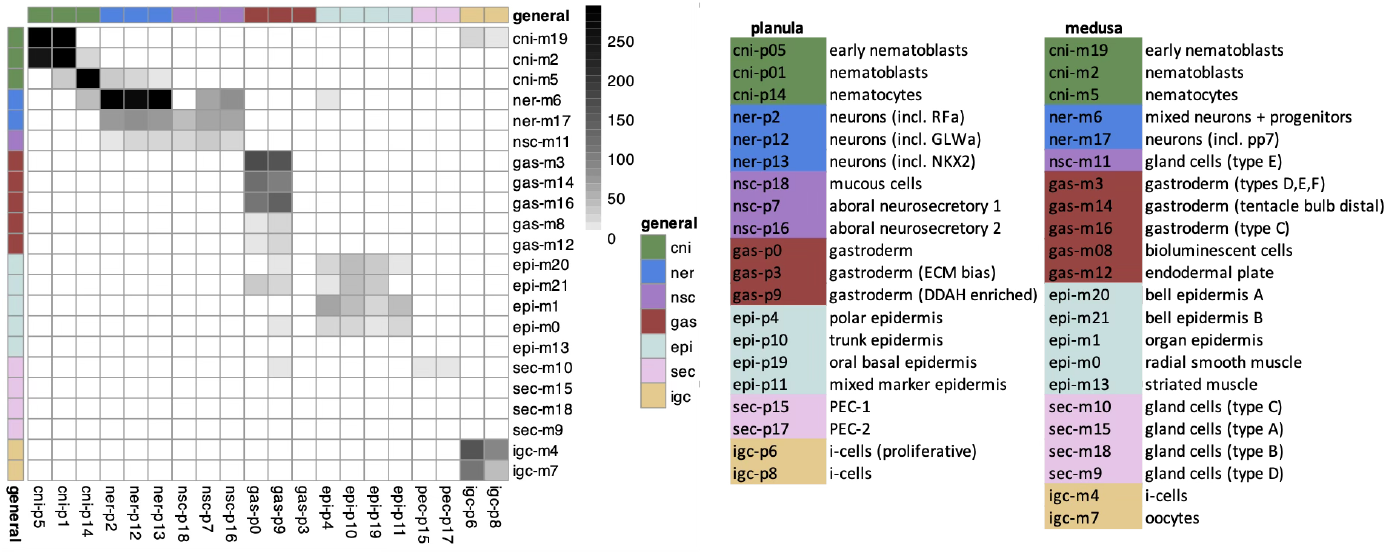
Significant numbers of shared marker genes between planula (x axis) and medusa (y axis) cell clusters. The colour ramp scale represents −log_10_(P) values of Fisher’s exact test for enrichment of shared marker genes (see methods). The general class of each cell is given by the color annotation (consistent with **Fig. 3**) with abbreviations as for **Fig. 1**.

**Figure 3:**
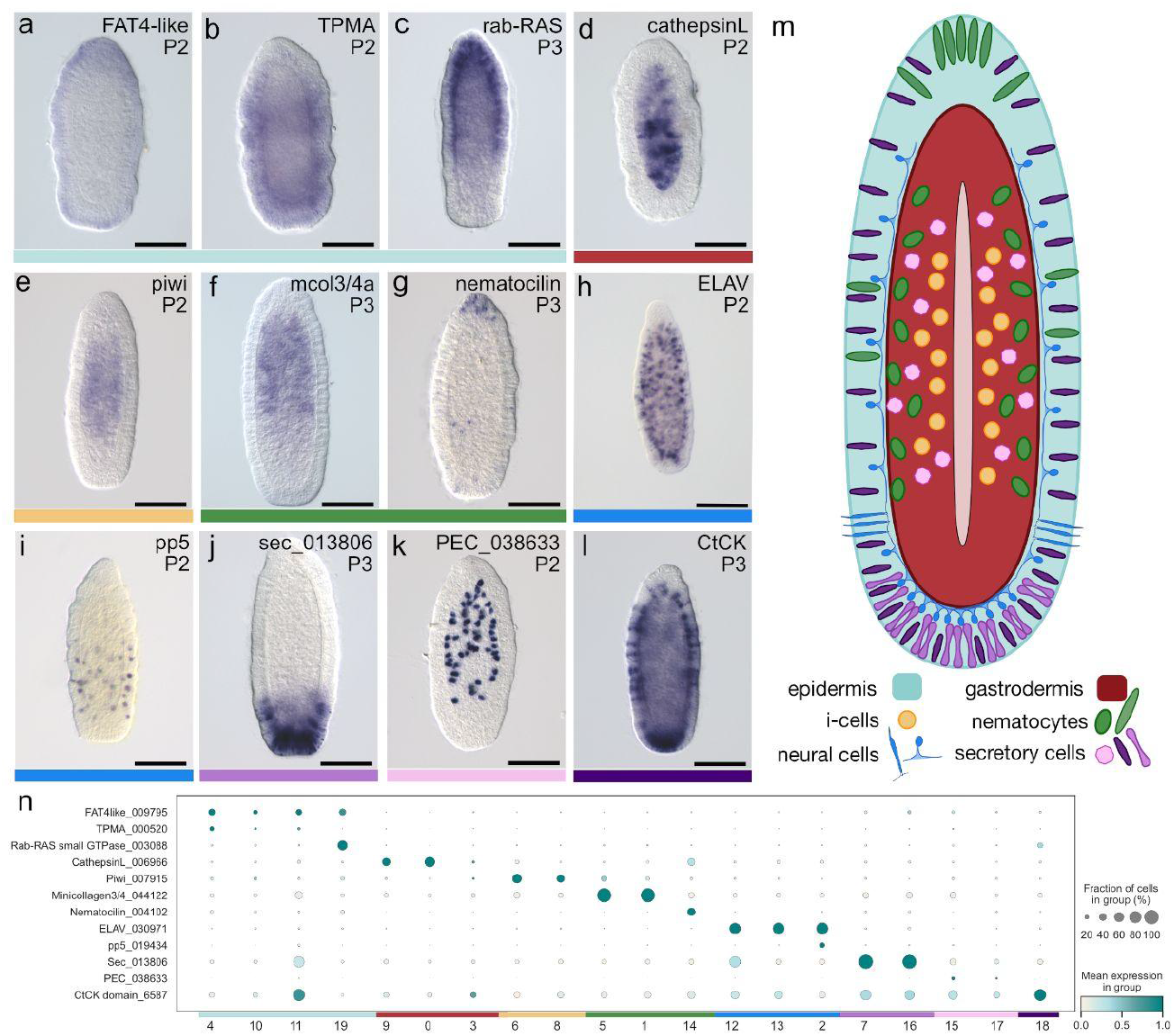
Examples of the distribution of major planula cell types. **a-l)** marker gene ISH; **m)** summary diagram summarising cell type distributions; **n)** Dot plot of marker gene expression across clusters. Cell type classes are colour-coded across panels. Planulae are oriented with the oral pole at the top.

**Figure 4:**
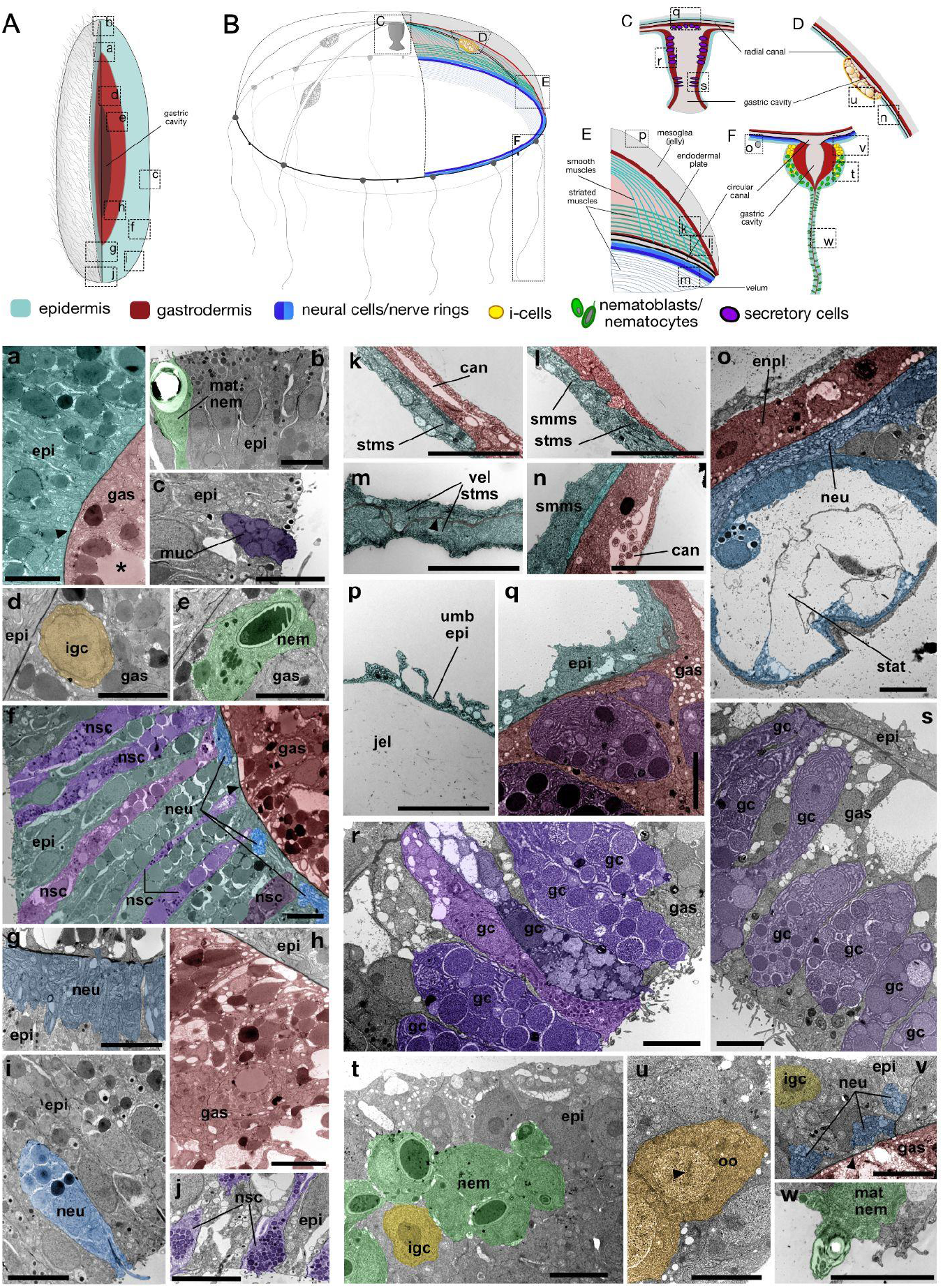
Ultrastructure of planula and medusa cells. **A)** Diagram of the planula (aboral end down). **B)** Diagram of the medusa. **C)** Medusa manubrium. **D)** Medusa gonad. **E)** Section of umbrella and velum. **F)** Tentacle bulb and tentacle. **a-w)** TEM micrographs illustrating ultrastructural features of planula **(a-j)** and medusa **(k-w)** cell types, from sites indicated by boxes on the diagrams. Planulae micrographs were taken from longitudinal sections. Colour coding of broad cell categories as in Figures 2 and 3. **a)** Basal region of the epidermis and gastrodermis layers, separated by mesoglea (black arrowhead), taken from near the oral pole. Most cells are epitheliomuscular cells, whose basal muscle fibres mingle with neurites alongside the mesoglea. Large round inclusions (asterisk) are likely lipidic yolk inherited from the egg. **b)** Oral region of the epidermis, including a mature nematocyte. **c)** Epidermis region including part of a mucous cell. **d)** Basal region of the epidermis and gastrodermis taken from the trunk region, with a putative i-cell in the gastrodermis. **e)** Similar region including a nematoblast in the gastrodermis. **f)** Overview of the epidermis and gastrodermis separated by a layer of mesoglea (black arrowhead) taken at the aboral pole. Neurosecretory cells are embedded between the epidermal epitheliomuscular cells. Neurites from sensory and ganglionic cells intermingle with their basal muscle fibres. **g)** Neural plexus at the aboral pole. Neurons are embedded in the epidermis layer and are adjacent to the mesoglea. **h)** Overview of the oral gastrodermis. **i)** Neurosecretory cells containing small electron-dense granules in the aboral/lateral epidermis. The presence of the apical cilium suggests a potential sensory function. **j)** Neurosecretory cells within the aboral epidermis. **k)** Subumbrella section showing striated muscle layer and gastrodermal cells of the radial canal. **l)** Subumbrella section showing layered smooth and striated muscle cells, and the endodermal plate. **m)** Velum showing striated muscle cells separated by a layer of mesoglea (black arrowhead). **n)** Subumbrella region towards the manubrium, showing smooth muscles and flagella within the radial canal. **o)** Specialised neuro-epidermal cells forming a statocyst adjacent to neurites of the nerve ring. **p)** Exumbrella epidermis. **q)** Basal region of the manubrium. The epidermis and the gastrodermis are separated by a layer of mesoglea. The gastrodermis includes several types of gland cells shown in **(r)** and **(s). t)** Epidermis of the tentacle bulb overlying nematoblasts and i-cells. **u)** Early stage oocytes in the gonad, showing typical condensed, paired chromosomes (black arrowhead). **v)** Epidermis of the tentacle bulb, including an i-cell and three neurite bundles of the nerve rings. The epidermis and gastrodermis are separated by a layer of mesoglea (black arrowhead). **w)** Mature nematocyte in the tentacle epidermis. Abbreviations for all panels: epi, epidermis; gas, gastrodermis; can, gastrodermal canal; nem, nematoblast; mat nem, mature nematocyte; muc, mucous cell; nsc, neurosecretory cell; igc, i cell/oocyte; enpl, endodermal plate; vel sms, velum striated muscle; smms, subumbrellar smooth muscle; stsm, sub-umbrellar striated muscle; umb epi, exumbrellar epidermis; neu, neurons; stat, statocyst; jel, jelly; gc, gland cells. Scale Bars: 5μm

An obvious correspondence between medusa and planula is seen with nematocytes which, at successive steps of differentiation, group in a series of cell clusters, with distinct expression patterns of key marker genes (medusa m19, m2, m5; planula p5, p1, p14), and ultrastructural similarity **(Fig. 4e, t)**. Immature nematocytes, termed nematoblasts, show strong expression of several minicollagens, components of the nematocyst, a pressurised capsule containing toxins [13]. In the planula, these are positioned in the gastrodermal region (**Fig. 3F, 3M; Fig. 4e)**. m19, representing the earliest stage nematoblasts in the medusa, also shows affinity with clusters p6 and p8 annotated as i-cells (**Fig. 2**), reflecting that most i-cells in the planula are destined for nematogenesis [14]. The differentiated nematocytes, in planula (p14) as in medusa (m5), express nematocilin **(Fig. 3N)**, whirlin and sans, bearing a gene expression signature reminiscent of mechanosensory hair cells [7]. In the planula, they are located in the ectoderm, clustered at the oral pole (**Fig. 3G** [14, 15]. Like medusa nematocytes and neurons, they also express the transcription factor POU4. The neuron clusters (medusa m6, m17; planula p2, p12, p13) are identifiable through common cell type-specific markers such as synaptotagmins, calmodulins and ELAVs (**Fig. 3h, S2C)**. In situ hybridisation with ELAV probes (**Fig. 3h**), and TEM (**Fig. 4g**) confirm the presence of neuronal projections close to the mesogleal layer, with a concentration in the aboral neural plexus (**Fig. 4g**). Sensory neuron bodies are interspersed between epidermal cells, and ganglionic cell bodies close to the mesoglea, with neuronal sub-populations distinguishable by their expression of specific marker genes such as the RFamide neuropeptide precursor (PP5) **(Fig. 3H,I; Fig. 4f,g,i**) [12]. The progression of cells through the nematogenesis pathway during planula development, along with other neurogenesis pathways, are considered more fully elsewhere [14].

Two broad groupings constitute the gastrodermis and epidermis. Epidermal cells were identified in 4 clusters (planula p4, p10, p11, p19), all expressing the cadherin Fat4-like (**Fig. 3A, 3N**). One of the clusters (p11) was characterized by non-specific expression of many highly expressed markers from other cell types, which we interpret as likely contamination from ambient mRNAs. Two others (p4 and p10) expressed genes related to muscle fibres such as Tropomyosin A (TPM-A, **Fig. 3A, 3N**) and showed enrichment of genes (such as radial spoke protein homologs, tektins and CFAPs, and GO terms associated with cilia (**Tab. S1**), indicating that these are ciliated cnidarian epitheliomuscular cells of the epidermal layer. Cluster 19 expresses such genes relatively poorly but instead is associated with many genes of the secretory machinery, such as RAB-Ras (XLOC_003088; Fig. 3C, 3N) an Arf (XLOC_005960), BLOC1S6 (XLOC_000404) (**Tab. S1**), and a fibrillar collagen (XLOC_042930). These cells were detected by ISH in a characteristic basal layer in the oral half of 3 day old planulae and likely contribute to establishing the mesoglea (**Fig. 3C**).

We identify 3 planula cell clusters as gastrodermis (p0, p3, p9), all expressing enzymes involved in intracellular digestion such as CathepsinL, as well as InnexinA and the previously identified *Clytic*-specific WegIE2 (**Fig. 5**). One cluster (p3) is enriched in proteins such as cadherins and collagens, as well as intracellular spectrin, associated with the establishment of epithelial layers and generating a basal extracellular matrix. The second main gastrodermal cluster (p0) additionally shows expression of FoxC, expressed in gastroderm cells in *Clytia* from the onset of ingression [16] and showing expression consistent with early differentiation of basal gastrodermal cells adjacent to the mesoglea, which have a characteristic vacuolated appearance (**Fig. S2a; Fig. 4f,h)**. p3 also shares some markers with the i-cell clusters, such as Piwi, Vasa and Nanos1, so potentially includes cells that have inherited germ-plasm-like aggregates of maternal transcripts of these genes, originally located at the egg animal pole [17]. The third gastrodermal cluster (p9) is smaller and is specifically enriched for DDAH (dimethylarginine dimethylaminohydrolase) transcripts.

**Figure 5:**
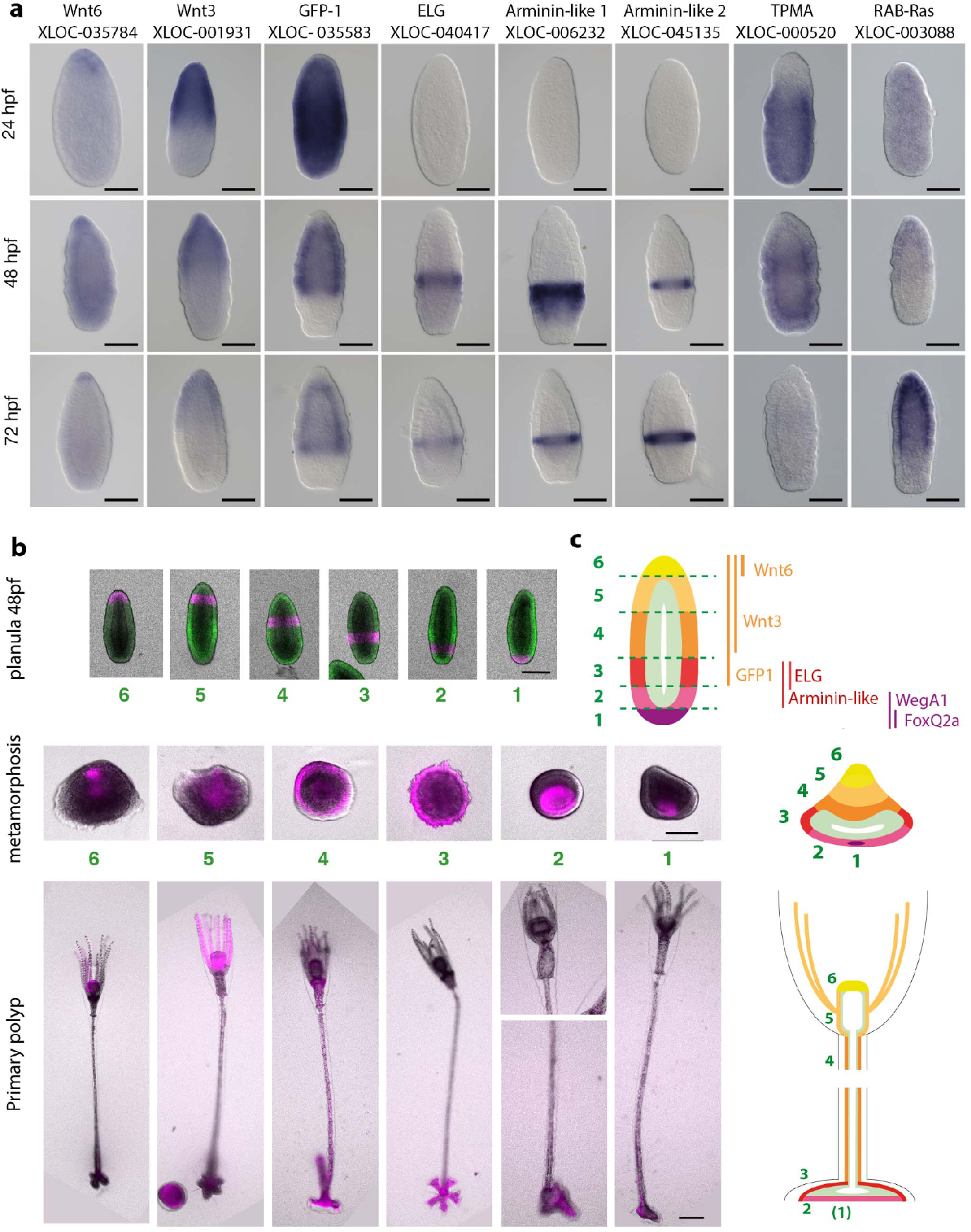
Ectoderm regionalisation along the oral-aboral axis predicts post-metamorphosis fates. **a)** In situ hybridisation for genes expressed in different zones at three stages of planula development as indicated. **b**,**c)** Fate mapping of six ectodermal zones defined along the aboral-oral axis by photoconversion of Dendra2 protein expressed from mRNA injected before fertilisation. **b)** Bright field images of planulae before (top row) and during (middle row) metamorphosis, and in primary polyps (bottom row). The fluorescence signal overlaid in green in the planula images corresponds to endogenous GFP1 and non-converted Dendra2, and all magenta overlays to photoconverted Dendra2. Photoconverted gastroderm cells mainly become interspersed during this period and are not distinguished here. c) Cartoons showing the mapping of planula epidermis regions to structures of the primary polyp. Oral is at the top in all images. Scale bars 100μm.

Two planula cell clusters, p6 and p8, shared markers with medusa i-cell and oocyte clusters, including Nanos1 and Vasa and were designated i-cells (defined broadly to include putative stem cells and early differentiating derivative cell types). Piwi expression was also enriched in these cells (**Fig. 3e, 3n)**. The larger cluster, p6, showed enhanced expression of the proliferation regulator PCNA (XLOC_031923). Znf845 (XLOC_017841), expressed early on the nematogenesis in *Clytia*, was enhanced in both p6 and p8, and the neurogenesis transcription factor Neurogenin in small subpopulations of each of their cells. By TEM, cells with stem cell-like features such as high nuclear-cytoplasmic ratio were observed in the gastroderm region, mostly containing nascent nematocysts (**Fig. 4d,e**). These observations are consistent with most ‘i-cells’ in the planula being engaged on the pathway to nematoblast production [14].

Two cell clusters (p7, p16), which we have termed neurosecretory [12], showed high expression of novel secreted proteins and were affiliated with neurons (**Fig. 2d**), but not typical neuronal markers such as ELAV[12] are distinguished ultrastructurally by their abundant large secretory granules **(Fig. 4f,j;** [18, 19]. A third cluster (p18), in turn showing similarities to these neurosecretory cells, was assigned as mucous secreting on the basis of shared markers with medusa mucous producing cells, and their characteristic morphology and distribution in the ectoderm along the entire axis of the planula (**Fig. 3L, 3N, 4c**, [15]).

The planula neurosecretory p16 cells were remarkable for extremely high expression of several genes coding for members of a family of homologous short secreted *Clytia*-specific proteins, which we term PF1 [12]. These proteins do not show sequence homology to any secreted proteins in other species and all share a structure characterized by conserved cysteines and the same 3D fold. Analysis of their expression across the *Clytia* life cycle using bulk transcriptome data [9] revealed that PF1s are expressed almost exclusively at the planula stage, with a maximum expression 2-3 days post fertilization, the period when larvae become competent to settle. These PF1 proteins are distinct from another group of short secreted proteins with conserved cysteines, which includes the ShK neurotoxins produced by *Nematostella*’s cnidocytes [20–22] as well as antimicrobial peptides (AMPs) produced by other cnidarians like damicornin and AmAMP1 in corals [23, 24] and aurelin in the jellyfish Aurelia [25].

In a similar vein, the neurosecretory p7 cells express the secreted protein XLOC_042849 at extremely high levels. Unlike PF1, which does not co-occur with known domains, the domain encoded by this gene co-occurs with Astacin-like proteases. Using Foldseek/Prost5 [26] we were able to detect distant similarity of *Clytia* family members to β-prism folds associated with membrane targeting [27].

Two further clusters (p15, p17) with pronounced similarity to each other showed no obvious neuronal signatures, but high expression of proteins encoding signal peptides. The transcriptome comparisons (**Fig. 1d, 2**) indicated a loose affiliation with other secretory cell types. *In situ* patterns of marker genes show a sparse and scattered distribution although with some bias towards the oral end of the larva (**Fig. S2 KK-l Z**). The expression profiles of markers for the two clusters altered over time, with some cells migrating to the ectoderm in day 3 planulae (P3) before becoming less abundant in older planulae (P4 in **Fig. S2**). These expression profiles are reminiscent of the distribution of the ‘granular cells’ described by Bodo and Bouillon [15], who speculatively proposed a role in the excretion of yolk breakdown products. These cells express several CTHRC (collagen triple helix repeat containing) genes, present in multiple copies in cnidarians and potentially involved in regulating cell migration [28]. We further noted expression of a number of lipoxygenases which oxidize polyunsaturated fatty acids to hydroperoxides, and the presence of specific Cytochrome p450s phylogenetically related to plant allene oxide synthases, which in animals are restricted to Cnidaria, *Trichoplax* and amphioxus [29]. Together these two enzymes are the first steps of eicosanoid biosynthesis. Eicosanoids in Cnidaria have been implicated in metamorphosis and neurotransmission, but little is really known [30, 31]. We refer to these as PEC cells — Putative Eicosanocytes. Comparing the lists of PEC cell marker genes to bulk RNA-seq data (see methods) shows striking overlap with bulk data that is likely to include theca - that is, stolon, polyp head and gonozooid (**Fig. S3**), so we suggest that these cells may be involved post-larval theca production.

### Regionalisation of the planula ciliated ectoderm

Previous work [16, 32, 33] has shown oral-aboral (O-A) regionalisation of the *Clytia* planula, including Wnt3 and its Frizzled receptors (Fz1, Fz3). We were able to detect evidence for this at the cellular level by sub-clustering ectodermal cell types (**Fig. S4**).

*In situ* hybridization of a panel of genes expressed in the epitheliomuscular cells of the ciliated ectodermal epithelial clusters p4 and p10 revealed a striking range of patterns defining distinct territories along the O-A axis (**Fig. 5;** see also [16, 33]). Genes expressed at the two poles of the planula (e.g. oral third: Wnt2 XLOC_031686, Wnt3 XLOC_001931, Wnt6 XLOC_035748, Bra and Bra2; aboral third: WegA1 XLOC_010822, ZnfA XLOC_014003) mapped predominantly within cluster p4, and those from central ‘trunk’ region, notably the GFP1 genes **(Fig. 5a**) within p10. From the expression patterns of such genes we designated six zones from aboral to oral along the planula and tracked the fates of epidermal cells in the primary polyp following metamorphosis using the photoconvertible fluorescent protein DENDRA-2 [34]. This enabled us to equate the regionalisation of ectodermal gene expression with specific fates, with discrete mapping indicating little or no more ectodermal cell mixing during metamorphosis. The oral tip zone 6 became the hypostome (mouth) of the primary polyp, zone 5 produced mainly tentacles, zone 4 the stalk, zone 3 the upper side of the basal disk and zone 2 the underside of the basal disk, while most of zone 1, enriched in the aboral neurosecretory cells likely discharged during settlement [12], was lost.

### Concordance of planula and medusa cell types

Inspection of the cell clusters showing significant similarity of marker genes between planula and medusa (**Fig. 3**) shows that there is not a one-to-one correspondence between planula and medusa. A proportion of this is likely to be due to minor differences in cluster compositions and technical aspects of cluster resolution. Equally however, correspondences and lack of correspondences can highlight important biological similarities and differences. Evolutionary and developmental considerations suggest that hierarchical models of cell similarity may be appropriate. To investigate this further, we inferred a tree-like structure of cell-type similarity using a Poisson based model of marker presence/absence implemented in the program biphy [35] (see methods and **Fig. 6**).

**Figure 6:**
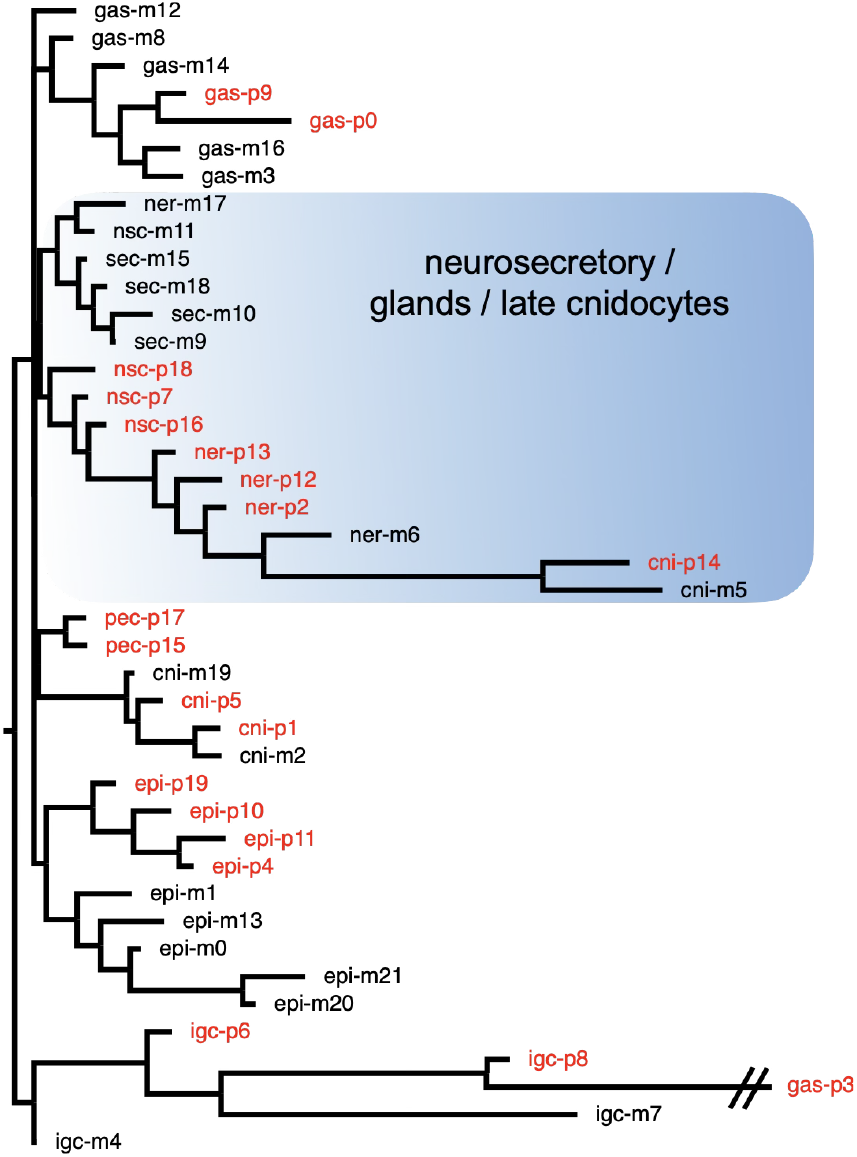
Tree based on shared marker gene presence or absence under a Poisson model (see methods), for medusa (black) and planula (red) cell types. Abbreviations: cni, nematocytes; epi, epidermis; gas, gastrodermis; gla, gland; igc i-cells/germ cells; ner, neuron; nsc, neurosecretory; pec, putative eicosanocyte; sec, secretory. The branch to gas-p3 is truncated.

In general, this analysis showed correspondences between groups of cells from planula and medusa, rather than precise identity between pairs of cells. Branches corresponding to gastroderm, ectoderm, nematoblasts and neurons contained both planula and medusa cell types. Terminal nematocytes of both planula and medusa branched within neurons as expected [7].

The epidermal cell types of the medusa are morphologically diverse (**Fig. 4k, l, m, n, p, q, t, v**), including striated (m13) and smooth muscle (m0) types as well as the exumbrellar ectoderm (m20 and m21) and epitheliomuscular cell layers encasing the gonad and manubrium (m1). These form a single clade with the planula epidermis clusters, within which the clusters from each stage are separate, indicating independent specialisation. Similarly, gastroderm clusters from medusa and planula form a clade, with p0 and p19 showing affinity with the medusa digestive gastroderm clusters m3, m14 and m16. Gastroderm cluster p3 falls outside this clade, its affinity with i-cells and oocytes in this analysis likely due at least in part to enrichment in stem cell gene mRNAs (see above).

Certain clusters we have annotated as gland cells in medusa (m9, m10, m15, m18) have no apparent planula counterparts. They are characterized by high expression of secreted enzymes such as trypsins and chitinases and are likely involved in extracellular digestion [7]. As the *Clytia* planula does not feed, their absence is simply explained. It is noteworthy though, that these gland cells do cluster within an extended neuro/secretory family of planula and medusa cells in our biphy based classification scheme, consistent with a common evolutionary origin [36], (**Fig. 6**), and show morphological similarity **(Fig. 4f,r,s)**. Medusa oocytes **(Fig. 4u)**, although related to i-cells, express specific markers related to the entry into meiosis, and are also not represented in our planula data; neither is the striated muscle type, although the striated muscle tropomyosin (XLOC_042542) is found enriched in one of the clusters we have assigned as putative eicosanocytes (p17), but without other muscle marker genes.

At the level of resolution presented here, there are three categories of strictly defined neurons (expressing ELAV and synaptotagmin) in the planula, corresponding to one in the medusa (p2,p12,p13 vs m6 (**Fig. 6**)). Two main basic types of cnidarian neurons, sensory and ganglionic, have been identified morphologically in the planula and are distributed in concentric zones around the aboral pole (**Fig. 4**) [12, 18]. Reanalysis of the scRNAseq neuronal clusters at higher resolution, shows that planula and medusa sub-populations can be distinguished with, for instance, different neuropeptide or other neurotransmitter complements. Thus, planula sensory neurons carry one of at least two distinct molecular signatures (expressing GLWamide versus HFRIamide neuropeptide precursors), but ganglionic neurons expressing precursors of RFamide populate mainly the trunk region and those expressing precursors of QYFamide the aboral neural plexus [7, 12]. Pairing these high-resolution complements between planula and medusa is highly dependent upon the precise levels of resolution of Leiden clustering (and thus the number of cell-type clusters identified), or the molecular features that are prioritized e.g. neurotransmitter identity vs. other molecules.

## Discussion

In common usage, cell-type definitions operate at differing levels of granularity (from as broad as neural to as specific as MB-RN-Spp1-Negative neurons [37]), softening the boundaries that can be applied to cell-type uniqueness [38]. Our single cell analysis of the *Clytia hemisphaerica* planula showed a similar complement of broad cell classes to the medusa, with both including neurons, nematocytes, secretory, epidermal, gastrodermal and interstitial (i.e. i-) cells. At a more fine grained level though, there are substantial differences in the cell complements.

At the broadest level the secretory cell category (**Fig. 6**) includes medusa digestive gland cells along with planula mucous cells (p18) and two neurosecretory cell types (p16, p7) positioned at the aboral pole [12]. Marker gene overlap showed that these three planula neurosecretory types showed transcriptome similarity with one of the medusa digestive gland cell types (m11; type E [7]), as well as neural cell types (p13, p12, p2 / m6, m17), suggesting neuro-secretory cells with common features are present in both stages. In contrast, the specialised medusa gland cell types characterised previously (m9, m15, m18; A,B and D of [7]) express specific sets of marker genes likely involved in extracellular digestion. We have addressed the issue of specialized cells involved in initiating the larval settlement response elsewhere [12], but note here that the neural and secretory cells we implicate do not appear to have direct counterparts in medusa, as judged by neurotransmitter use (e.g. HFRIamide/TauD neurons) and other secreted proteins.

Within the epidermal cell category, it is notable that the genes expressed in planula cells show a strong enrichment of ontology terms for genes involved in cilia formation, as might be expected. Although we can detect significant similarity between these planula cells and medusa exumbrella cells, we do not find a comparable enrichment of cilia GO terms in medusa bell epidermis, consistent with these motile cilia being a feature of planulae. The several additional muscle and epidermal cell types of the medusa associated with particular organs (manubrium, gonad, tentacle) and functions (swimming, tentacle contraction, gamete expulsion etc) are highly specialised but nevertheless share transcriptional signature components (**Fig. 2 and 6**). Similarly the gastroderm tissues of the medusa are more complex than those of the planula, including a vascular system linking compartments in the manubrium, gonad and tentacle bulbs. Some medusa gastroderm cells are characterised by features shared extensively with the planula involved with nutrient uptake, intracellular digestion and storage, while others show distinct specialisations for instance associated with structural features such as the endodermal plate.

The *Clytia* planula and medusa are not contiguous stages. The planula first metamorphoses to a primary polyp, ultimately yielding a colony including specialised gonozooids that bud medusae. A tough extracellular theca surrounding all colony structures hampers the generation of cell suspensions and hence single cell transcriptomic data and as yet, none of the latter are available. Our lineage tracing experiments demonstrate continuity between epidermal regions of the planula and polyp **(Fig. 5)**. The degree to which this continuity manifests and the transcriptomic level is an obvious follow-up question. One of the more distinct planula cell populations, PECs (p15, p17) shows strong enrichment of genes found in bulk RNA-seq libraries of polyp structures with substantial theca components (stolon, gonozooid, polyp head; **Fig. S3**). These structures may have other cell-types in common besides those that secrete the theca though, such as types involved in immunity.

This work has illustrated that the diversity of cell identities observed between distinct life-cycle stages in *Clytia* generally reflect independent variations on a set of basic themes, specialised to meet the specific needs of lifestyles and morphologies.

## Methods

### *Clytia* medusae culture

*Clytia* female jellyfish (Z4B strain) were cultured in Kreisel tanks for approximately 10-14 days and fed twice a day as previously described (Lechable et al., 2020).

### Cell dissociation of *Clytia* medusae

Dissociation of medusae was carried out without the use of digestive enzymes. The new samples Medusa2 and Medusa3 were prepared from 10-14-day-old medusae, which had not yet reached sexual maturation, were transferred to a petri dish (ø = 100mm) filled with Calcium/Magnesium-free artificial sea water (Ca/Mg-free ASW: 530.5mM NaCl, 10.7 mM KCl, 3.5 mM NaHCO_3_, 11.3mM Na_2_SO_4_, pH=8) and placed into a 40μm cell mesh positioned within the petri dish. Medusae were washed three times in Ca/Mg-free ASW by serial transfer in petri dishes containing fresh Ca/Mg-free ASW to dilute the cations stepwise. Following the last wash, medusae were incubated in fresh Ca/Mg-free ASW for ten minutes. Following incubation, excess seawater was removed, and medusae were mechanically dissociated by gently pressing the tissue through the 40 μm mesh. Cells were recovered by washing the cell mesh with 500 μl of Low-Calcium/no-Magnesium artificial sea water (LowCa ASW: 460mM NaCl, 9.9mM KCl, 1.4 mM CaCl_2,_ 10mM HEPES, pH=7.6). Cell suspension was collected in a 2ml Lobind tube. 10μl of cell suspension was isolated for manual counting with a Neubauer improved counting chamber (Sigma-Aldrich BR717810-1EA) to estimate cell concentration. Unlike the dataset Medusa1 documented in our previous ‘WHAM-Seq’ study involving sample multiplexing, these Medusa2 and Medusa3 samples were loaded and encapsulated without prior fixation.

### *Clytia* planulae culture

Fertilisation was carried out by mixing spawned gametes from laboratory-cultured adult medusae (Z23 female strain and Z4C male strain). Embryos and larvae were cultured in filtered seawater (Red Sea Salt brand) with 1:25000 penicillin and streptomycin solution at 17°C [39].

### Cell dissociation and fixation of *Clytia* planulae

The cell dissociation protocol for *Clytia* planulae was based on initial trials comparing data obtained using unfixed dissociated cells and ones fixed in different ways. Dissociated cells for the Planula-3 and Planula-4 samples were prepared as previously described [12] (**Tab. S3)**. Planula-1 and Planula-2 samples **(Tab. S3)** were collected at about 50 hours post fertilisation (hpf) in a 40μm cell mesh placed in a petri dish filled with Ca/Mg-free ASW and washed three times by serial transfer into fresh Ca/Mg-free ASW, as described above for the medusa. Cell dissociation was carried out by incubating the planulae in Ca/Mg-free ASW containing 40U/mL SUPERase•In™ RNase Inhibitor (20U/mL, Invitrogen) for ten minutes. Excess seawater was removed from the strainer, and planulae were gently pressed against the mesh. Cells were recovered by washing the cell mesh with 500 μl LowCa ASWcontaining 40U/mL SUPERase•In™ RNase Inhibitor. Cell concentration was estimated by isolating 10μL of cell suspension and staining with Hoechst 33258 (bisBenzamide, Sigma-Aldrich; 1:1000 of a 1 mg/ml stock in water). Cells were counted manually with a Neubauer improved counting chamber (Sigma-Aldrich BR717810-1EA). Dissociated cells were immediately fixed with 80% ice-cold methanol (Planula-1 sample) as in [7] or with ice-cold ACME solution (Planula-2 sample) [40] and 40U/ml SUPERase•In™ RNase Inhibitor (20 U/μL), then incubated for 30 min at −20°C or on ice.. Following incubation, a solution of Phosphate Buffered Saline (PBS: 110 mM NaCl, 1.9 mM KCl, 8mM Na_2_HPO_4_, 2 mM KH_2_PO_4_ pH 7.2)-1%BSA and 40U/ml SUPERase•In™ RNase Inhibitor (20 U/μL) was added to the cell suspension in a 1:1 ratio and mixed by gently pipetting to resuspend the cells. Cells were pelleted by centrifugation for 10 min at 700 rcf in a swinging bucket centrifuge. Superanatant was discarded, and cell pellets were resuspended with 800μL of 1%BSA in PBS containing 100U/ml SUPERase•In™ RNase Inhibitor (20 U/μL). Finally, DMSO was added to the cells to a final concentration of 10% followed by snap-freezing in liquid nitrogen and storage at −80°C until encapsulation. Aliquots of dissociated cells were prepared over several days.

### Library preparation and sequencing

All planula and medusa cell samples (i.e. without regard to dissociation or fixation method) were encapsulated using the 10X Genomics Chromium system and cDNA libraries were prepared according to the Chromium Next GEM Single Cell 3’ library preparation protocol v3.1.

For the medusa samples, freshly dissociated cells were immediately mixed with the 10X Master Mix, the solution was added to a pre-loaded 10X microfluidic chip, and the encapsulation reaction was started. Cell encapsulation targeting 10000 cells, cDNA library preparation according to the Chromium Next GEM Single Cell 3’ library preparation protocol v3.1 and sequencing of single cell libraries (NextSeq 500/550 Midouput and NextSeq 500 Midoutput kit, 75 cycles kits) were performed by the UCAGenomiX platform, partner of France Génomique, at the Institut de Pharmacologie Moléculaire et Cellulaire (IPMC, Sophia Antipolis, Nice, France) and at the ‘GeneCore’ Genomics facility at the European Molecular Biology Laboratory (EMBL Heidelberg, Germany).

Fixed planula cells were shipped to the Centre for Genomic Regulation (CRG, Barcelona, Spain; samples Planula-1 and Planula-2) where preliminary steps to encapsulation were performed, or processed at the UCAGenomiX platform, partner of France Génomique, at the Institut de Pharmacologie Moléculaire et Cellulaire (IPMC, Sophia Antipolis, Nice, France; Planula-3 and Planula-4) as previously described [12]. All samples included in the final datasets are listed in **Fig. S3**. Samples were thawed on ice, and tubes were centrifuged at 1000xg in a swinging bucket centrifuge. The supernatant was discarded, and the cell pellet was washed with 0.5% BSA in PBS containing 40 U/mL RNase inhibitor. Cells were washed twice to eliminate the DMSO. Aliquots were combined, and the cells were finally transferred to Low Binding tubes in a total volume of 500μL PBS-0.5%BSA with 100U/mL RNase inhibitor.

A sorting step was performed for Planula-1 and Planula-2 samples prior to encapsulation to eliminate any ambient RNA and debris,, to estimate the cell concentration of Planula 1 and Planula 2 samples after thawing and washing, 6 μL of cell suspension were stained with 1:1 PBS-1:100 DAPI and counted using a Countess machine (Invitrogen). Single cell target populations (mainly 2n and 4n DNA content) were purified from debris and aggregates using a BD Aria Fluorescence-Activated Cell Sorter (FACS) and encapsulated using the 10x Genomics platform, as previously described (https://doi.org/10.1186/s13059-021-02302-5). Briefly, cells were thawed and washed in PBS containing 0.5% BSA and RNase inhibitor (100U/mL), adjusted to 1-1.5M cells/mL, and stained for FACS using the nuclear dye DRAQ5 (1/300 dilution from a 5 mM stock) and the cytoplasmic dye Concanavalin-A-Alexa Fluor 488 (4μL of 1mg/mL stock). Between 7000 and 10000 single cells were sorted directly into Master Mix lacking RT enzyme C (single cell 3’ reagents kit v3.1) using a 100-μm nozzle and single-cell purity mode. Following addition of RT enzyme C, cells were immediately loaded onto 10x Genomics Chromium chips, and single-cell 3’ GE libraries were constructed according to manufacturer’s protocol.. For these samples, cell encapsulation targeting 4000 to 6000 cells, cDNA library preparation and sequencing of single cell libraries (NextSeq 150 Midoutput and NextSeq 75 High output kits) were carried out by the Genomics facility at the Centre for Genomics Regulation (CRG, Barcelona, Spain).

### Processing of single-cell transcriptome data

Analysis python notebooks are available at: http://github.com/rcply/planula_vs_medusa. In brief, reads for each library were mapped to a chromosome-level *Clytia hemisphaerica* genome and gene models [12] using STARsolo (v2.7.9a) [41]. Output count matrices were analyzed separately for medusa and planula libraries using Scanpy (v1.11) [10], with minima of 100 genes per cell and 3 cells per gene, and less than 5% mitochondrial reads for medusa; and 200 genes per cell with less than 10% mitochondrial reads for planula. Within each life stage (planula, medusa) we integrated libraries using Harmony [11], identifying 75 neighbors using the integrated PCA and then Leiden clustering at 0.5 resolution. Planula and medusa were each processed computationally as 3 library batches (details in **Fig. S3**). Pseudobulk matrices were created for planula and medusa cell-types by aggregating the counts for each Leiden cluster within each batch, with the cluster number as the condition (all clusters were represented in all conditions), and the batch forming part of the differential expression analysis design. For each cell type, contrasts were performed against the combined set of all other cell types for that stage (planula or medusa) using DESeq2 [42]-see code for details.

### Shared marker genes

We took all significant marker genes for each cell cluster, and for each pair of cell clusters constructed a contingency table *((a,b),(c,d))* such that *a* was the number of marker genes shared by the two clusters, *b* and *c* the number of marker genes unique to each cluster of the pair, and *d* the number of marker genes remaining in the union of all other cell cluster markers. This table was used as the basis for Fisher’s exact test for each cell pair. We did three comparisons: planula vs. planula; medusa vs medusa and planula vs medusa. We took −log_10_ of the *P*-value for the pairwise comparison as the cell value in the heatmaps illustrating all pairwise comparisons.

### Gene ontology analysis

We used PANTHER family assignments [43] (v17.0) to retrieve GO term annotation for genes, and then used ‘find_enrichment.py’ from the GOATOOLS system to determine significantly overrepresented terms [44].

### Hierarchical analysis of marker gene content

In order to obtain a hierarchy of cell type similarity, we constructed a binary matrix of marker gene presence/absence across all cell type clusters (i.e. medusa and planula). We analysed the matrix using biphy, a method to infer phylogenies from binary character presence/absence data by modelling character gain and loss as a Poisson process (https://github.com/milliescient/biphy [35]), correcting for the inability to observe genes that were absent in both datasets. We merged the results of 10 independent outputs and obtained a consensus tree depicting a hierarchy of shared marker gene content between clusters across planula and medusa stages.

### *In situ* hybridisation

Larvae were fixed at 24, 48, 72 and 96 hpf in 3.7% formaldehyde, 0.2% glutaraldehyde in PBS on ice for 40 min. Specimens were then washed with PBS + 0.1%Tween 20(Sigma-Aldrich, PBST), dehydrated stepwise in methanol on ice, and stored in 100% methanol at −20°C. Colourimetric in situ hybridisation was performed as previously described for all stages [45] (or, in some cases, using a robot (Intavis AG, Bioanalytical Instruments) as described in [7]. Probes were generated by polymerase chain reaction (PCR) from cDNA clones of the *Clytia* expressed sequence tag (EST) collection [7, 46] or from planula cDNA. Oligonucleotide primers were designed with PrimerBlast (https://www.ncbi.nlm.nih.gov/tools/primer-blast/) and cloned in the pGemT-easy vector (Promega, CAT#A1360) according to the manufacturer’s recommendation. Sequences of the oligonucleotide primers and EST identification names are provided in **Tab. S2**.

### DENDRA2 lineage tracing

The coding sequence for Dendra2 [34] was adapted to the codon usage of *Clytia* [47]. A T7 promoter-Dendra2 ORF DNA template was obtained by PCR and mRNA was synthesised using mMessage mMachineTM T7 Transcription Kit (Ambion), followed by poly-A addition using the Poly(A) Tailing Kit (InVitroGen). Solutions of mRNA at 0.4 - 1.0μg/μl were injected into freshly spawned eggs prior to fertilisation using an Eppendorf nanoject injection system as described in detail previously [48]. Photoconversion was performed on a Leica Stellaris confocal microscope using the LasX software FRAP mode with a 10X objective. The larvae were anesthetized with MgCl_2_ and mounted temporarily between slides and coverslips separated by plasticine spacers. Each defined region was exposed to 30 seconds scanning of the 405 nm Laser beam, at 100% power.

Metamorphosis was then triggered by GLWamide treatment [39] and imaged on an Axiozoom microscope at successive times.

### Electron Microscopy

Planula at 2 days and 3 days after fertilisation, and female medusae fed daily in culture for 10-14 days after budding, were fixed using a sequential Osmium Tetroxide - Glutaraldehyde method [49]. After embedding in Epon resin, ultrathin sections were stained with lead citrate and imaged with a JEOL 1400 transmission electron microscope.

## Acknowledgments

We thank Tsuyoshi Momose (IMEV, Villefranche-sur-mer) for providing the DENDRA2 construct that he customised for use in *Clytia*; Bianka Baying and Vladimir Benes (Genecore, EMBL) for help generating early datasets. This work was funded by the H2020/Marie Skłodowska-Curie ITN “EvoCell” (grant agreement no. 766053) and the French ANR-21-CE13-0024 “MOPSEA”. Microscopy facilities were provided by the Imaging Platform and *Clytia* cultures were maintained by the Service Aquariologie of Centre de Ressources Biologiques Marines (IMEV – FR 3761) of Institut de la Mer de Villefranche, both supported by EMBRC-France (ANR-10-INBS-02). The UCA GenomiX Platform is supported by the Conseil Départemental des Alpes Maritimes (2016-294DGADSH-CV), and of the National Infrastructure France Génomique (Commissariat aux Grands Investissements) [ANR-10-INBS-09-03 and ANR-10-INBS-09-02]. Cell sorting trials with a BD FACAria III cell sorter (BD Biosciences) were performed at the “Microscopie Imagerie Cytométrie Azur” facility.

## Supplementary figures

**Figure S1:**
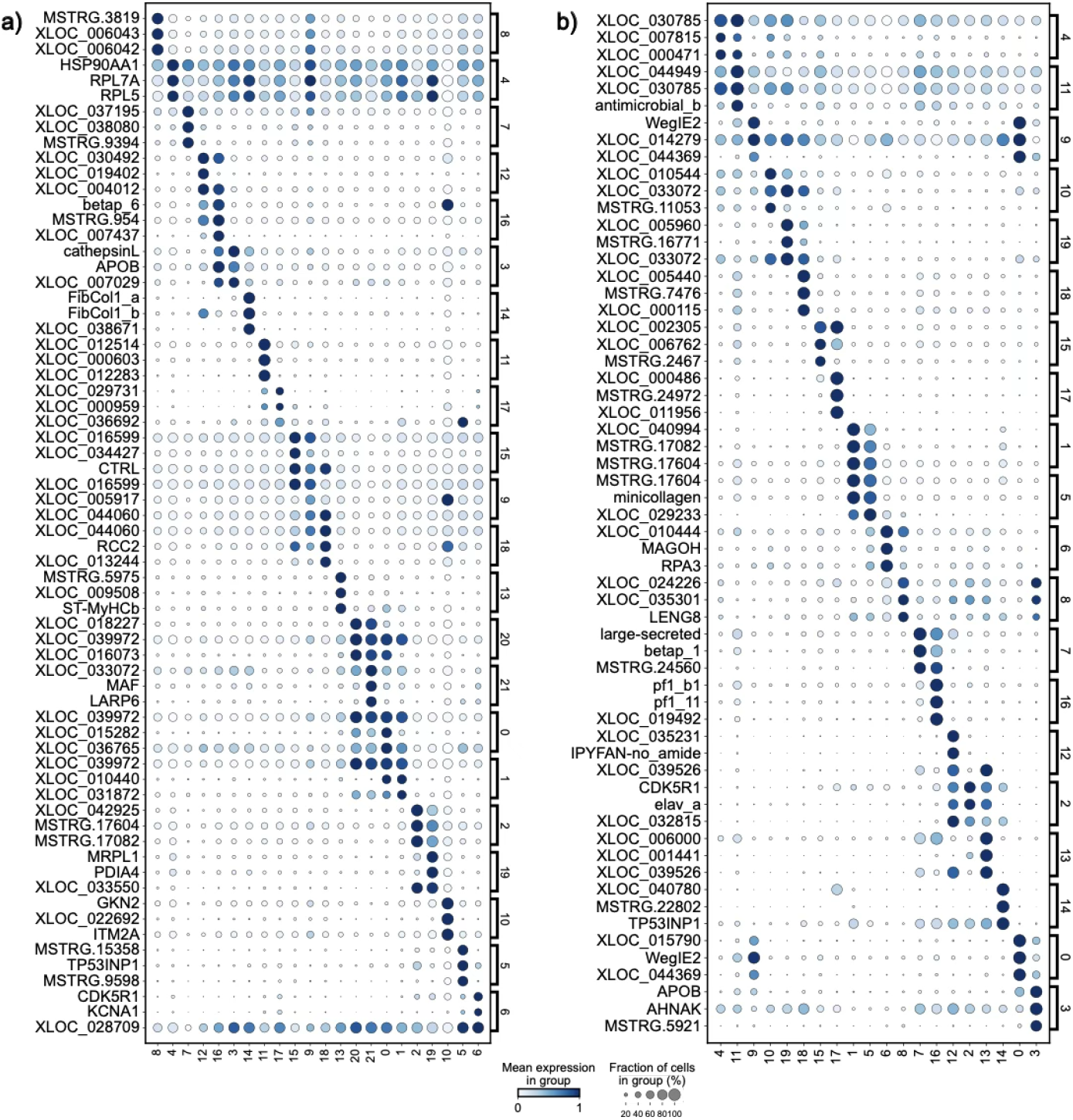
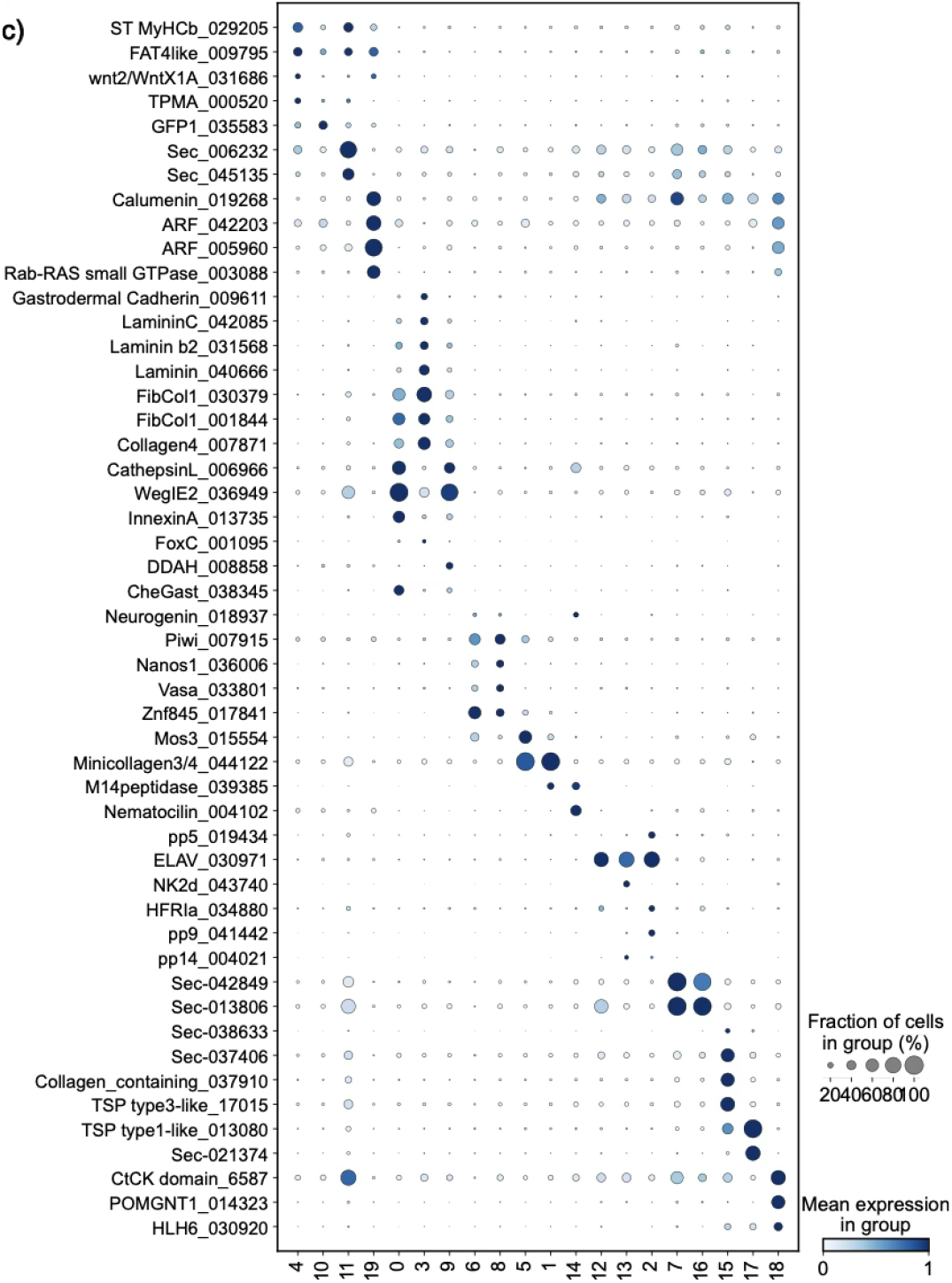
top 3 marker genes for each of the **a)** medusa and **b)** planula Leiden clusters.**c)** selected planula cell type markers discussed in the text, by Leiden cluster. Suffix numbers in gene names are database accessions.

**Figure S2.**
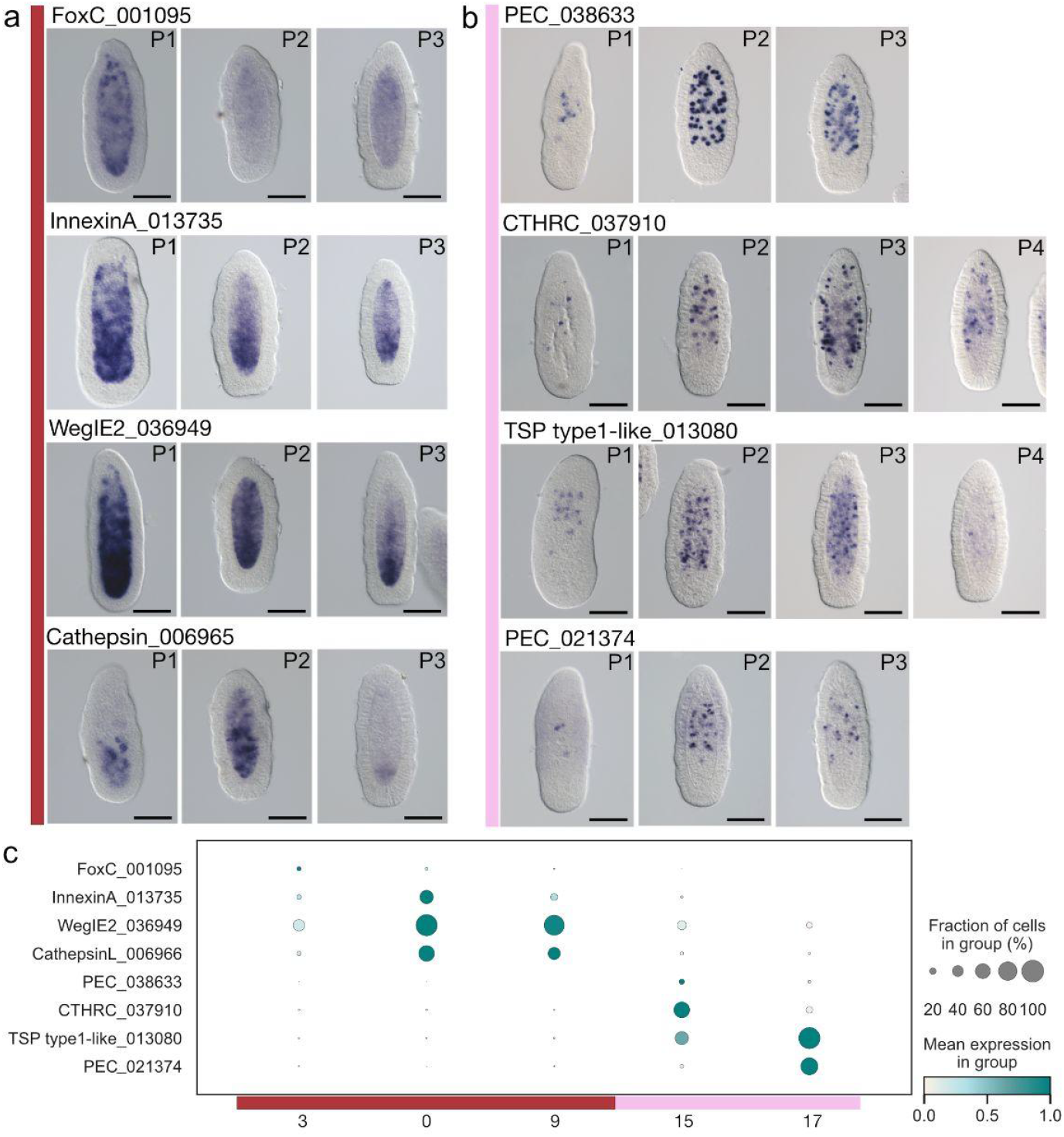
Gastroderm diversity in the two-day planula. **a)** In situ hybridisation for genes expressed in the gastroderm in 2-day planulae as indicated. **b)** In situ hybridisation for PEC cell markers. **c)** Dot plot comparing marker expression between clusters. Oral is at the top in all planula images. Scale bars 100μm

**Figure S3:**
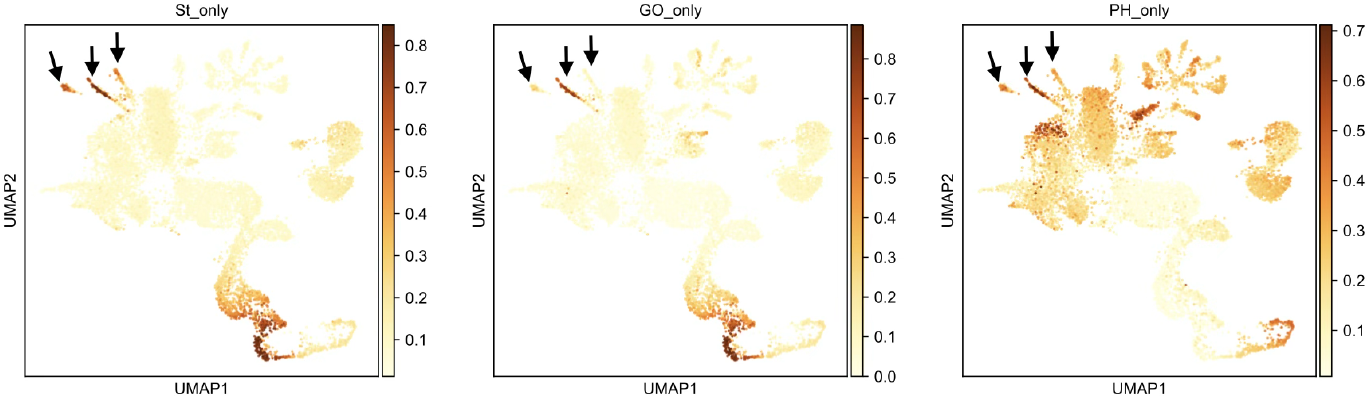
Planula PEC cell overlap with bulk RNA-seq data from theca containing samples. St bulk stolon; GO bulk gonozooid; PH bulk polyp head. Arrows point to PEC cell clusters.

**Figure S4:**
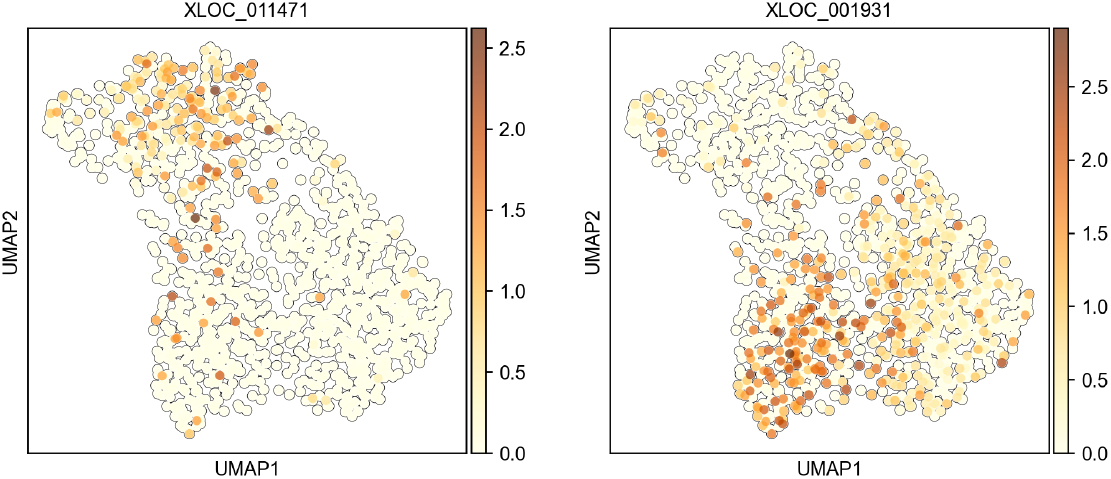
An Fz domain containing secreted protein (XLOC_011471, an aboral marker) and Wnt3 (XLOC_001931, an oral marker) show populations of epidermal cells (reclustered p4) with distinct regionalisation.

## Supplementary tables

**Supplementary table 1:** Enriched Gene Ontology terms for medusa and planula cell clusters.

**Supplementary table 2:** List of marker genes used in this study.

**Supplementary table 3:** New samples reported in this study (Planula 1, Planula 2, Medusa 2, Medusa 3). Planula 3 and Planula 4 are reported in Ramon-Mateu et al., 2025, and Medusa 1 is reported in Chari et al., 2021.

## Notes

### Competing Interest Statement

The authors have declared no competing interest.

